## Supplemental Table 1 for "Fish CDK2 recruits Dtx4 to degrade TBK1 through ubiquitination in the antiviral response"

SI Table. Primers used in this study.

| **Name** | **Sequence (5’→3’)** | **Purpose** |
| --- | --- | --- |
| pCMV-HA-CDK2-F | CCGGAATTCCGATGGAGTCTTTTCAGAAAGTG | Eukaryotic expression |
| pCMV-HA-CDK2-R | CCGCTCGAGTTATAGGCGTAAAGGAGGCAC |  |
| mCherry-MAVS-F | CGCGGATCCGCCACCATGTCACTGACACGTGAGC |  |
| mCherry-MAVS-R | CCCATCGATATGATTGAGCTTCCAG |  |
| mCherry-TBK1-F | CGCGGATCCGCCACCATGCAGAGTACGGCCAAT |  |
| mCherry-TBK1-R | CCGGAATTCCGCATCCGCTCCACTG |  |
| pEGFP-N3-CDK2-F | CTAGCTAGCATGGAGTCTTTTCAGAAAGTG |  |
| pEGFP-N3-CDK2-R | CGCGGATCCTAGGCGTAAAGGAGGCAC |  |
| pCMV-Myc-MAVS-F | CCGCTCGAGCGATGTCACTGACACGTGAGC |  |
| pCMV-Myc-MAVS-R | ATTTGCGGCCGCTTAATGATTGAGCTTCCAG |  |
| pCMV-Myc-TBK1-F | CGCGGATCCCGATGCAGAGTACGGCCAAT |  |
| pCMV-Myc-TBK1-R | AACTCGAGTCACATCCGCTCCACTG |  |
| pCMV-Tag2C-TBK1∆N-F | CCGGAATTCCGATGCAGAGTACGGCCAATTACCTGTGGATGATGTCCGACCTGACAGCGAACCTCTTC |  |
| pCMV-Tag2C-TBK1∆N-R | AACTCGAGTCACATCCGCTCCACTG |  |
| pCMV-Tag2C-TBK1∆C-F | CCGGAATTCCGCTGGGTCAGGGAGCCACAGC |  |
| pCMV-Tag2C-TBK1∆C-R | AACTCGAGTTAGTTGTAAGTGTGTATGTAG |  |
| pCMV-Tag2C/HA/Myc-Dtx4-F | CGGAATTCCGATGGACACGATGTTGCTG |  |
| pCMV-Tag2C/HA/Myc-Dtx4-R | CCGCTCGAGTCACAGGAGCTGTTCCTC |  |
| pCMV-Myc-trim11-F | CGGAATTCCGATGGCGTCCTCCAGTGTTTTG |  |
| pCMV-Myc-trim11-R | AAAAGCGGCCGCTTAATAGACAGGTGTGATG |  |
| pCMV-Myc-Traip-F | CGGAATTCCGATGCCCATTCGAGCATAC |  |
| pCMV-Myc-Traip-R | CCGCTCGAG TTATTCCAAGAACCCGTCC |  |
| pCMV-Myc-Socs3a-F | CGGAATTCCG ATGATAACCCACAGCAAG |  |
| pCMV-Myc-Socs3a-R | CCGCTCGAGTTAAATAGGGGCGTCATAC |  |
| pCMV-Myc-Dtx4∆WWE-F | CGGAATTCCGATGGACACGATGTTGCTGGCTGTCTCTGGGCCTCTGCCAAA | Eukaryotic expression |
| pCMV-Myc-Dtx4∆WWE-R | CCGCTCGAGTCACAGGAGCTGTTCCTC |  |
| pCMV-Myc-Dtx4∆RING-F | AGAAGGTCAGGAGTCCACCAGTAAAGACAGGCAACCAGCC |  |
| pCMV-Myc-Dtx4∆RING-R | GGCTGGTTGCCTGTCTTTACTGGTGGACTCCTGACCTTCT |  |
| pCMV-Myc-Dtx4∆DTC-F | CGGAATTCCGATGGACACGATGTTGCTG |  |
| pCMV-Myc-Dtx4∆DTC-R | CCGCTCGAGTTACTTTACTCCATATATAGT |  |
| sh*-cdk2*#1-F | CCGGGCACTGCCATACGTGAGATCTCTCGAGAGATCTCACG TATGGCAGTGCTTTTTG | Knock-down |
| sh-*cdk2*#1-R | AATTCAAAAAGCACTGCCATACGTGAGATCTCTCGAGAGATCTCACGTATGGCAGTGC |  |
| sh-*cdk2*#2-F | CCGGGGGTGGTTTATAAAGCCAAGACTCGAGTCTTGGCTTT ATAAACCACCCTTTTTG |  |
| sh-*cdk2*#2-R | AATTCAAAAAGGGTGGTTTATAAAGCCAAGACTCGAGTCTTGGCTTTATAAACCACCC |  |
| sh*-dtx4*#1-F | CCGGAGTACCAGTGAAGAATCTTAACTCGAGTTAAGATTCTTCACTGGTACTTTTTTG |  |
| sh-*dtx4*#1-R | AATTCAAAAAAGTACCAGTGAAGAATCTTAACTCGAGTTAAGATTCTTCACTGGTACT |  |
| sh*-dtx4*#2-F | CCGGGGATTTCCCCGTCATTGTTATCTCGAGATAACAATGACGGGGAAATCCTTTTTG |  |
| sh*-dtx4*#2-R | AATTCAAAAAGGATTTCCCCGTCATTGTTATCTCGAGATAACAATGACGGGGAAATCC |  |
| *Dr-cdk2-*F | CAGAAAGTGGAGAAGATCGGAG | Real-time PCR  Real-time PCR |
| *Dr-cdk2-*R | AGTCCATAAACCTCTTCAG |  |
| *Dr-β-actin*-F | CCGTGACATCAAGGAGAAGCT |  |
| *Dr-β-actin*-R | TCGTGGATACCGCAAGATTCC |  |
| *epc*-*β-actin-*F | CACTGTGCCCATCTACGAG |  |
| *epc*-*β-actin-*R | CCATCTCCTGCTCGAAGT |  |
| *epc*-*cdk2-*F | GTGTTCCCAGCACTGCCATA |  |
| *epc*-*cdk2-*R | GGTAACTCTTCACGAGTGGCA |  |
| *epc-ifn*-F | ATGAAAACTCAAATGTGGACGTA |  |
| *epc*-*ifn-*R | GATAGTTTCCACCCTTTCCTTAA |  |
| *epc*-*vig1-*F | AGCGAGGCTTACGACTTCTG |  |
| *epc*-*vig1-*R | GCACCAACTCTCCCAGAAAA |  |
| *p-*F | TTGGACCTGGGATAGTGA |  |
| *p-*R | CTTGCTTGGTTTGTGGG |  |
| *n-*F | TGAGTGCTGAGGACGAT |  |
| *n-*R | TTTGTGAGTTGCCGTTA |  |
| *g-*F | CGACCTGGATTAGACTTG |  |
| *g-*R | AATGTTCCGTTTCTCACT |  |
| *l-*F | GCCCACTTTGCATCCAGTCC |  |
| *l-*R | GAGATGCCACAGACTCCTCC |  |
| *m-*F | TACTCCTCCCACTTACGA |  |
| *m-*R | CAAGAGTCCGAGAAGGTC |  |
